# Bacterial exploration of solid/liquid interfaces: developing platforms to control the physico-chemical microenvironment

**DOI:** 10.1101/2025.07.25.666890

**Authors:** Mathieu Letrou, Kennedy Chagua Encarnacion, Rebecca Mathias, Yeraldinne Carrasco Salas, Sofia Gomez Ho, Elena Murillo Vilella, Lionel Bureau, Sigolène Lecuyer, Delphine Débarre

## Abstract

Bacterial long-term contamination of surfaces is a promiscuous phenomenon often linked to harmful processes. Early bacterial exploration of interfaces, governed by adhesion and individual motility, is a known determinant of the subsequent development and persistence of bacterial colonies. However, the mechanisms by which bacteria integrate various environmental signals at these interfaces and modulate their behavior in response remain poorly understood. Here we present methods for designing precisely controlled microenvironments that enable the manipulation of both physical and chemical properties of solid-liquid interfaces, and also permit *in situ* monitoring of bacteria at these interfaces within microfluidic flow cells. Our aim is to provide an innovative toolbox for the interdisciplinary research community focused on elucidating the complex processes underlying bacterial surface exploration. We illustrate its use here by examining the surface motility of the pathogen *Pseudomonas aeruginosa*.

## 1 Introduction

Many bacterial species alternate between a motile lifestyle, where individual cells swim freely, and a sessile state, where they attach to surfaces, proliferate into microcolonies, and eventually form biofilms [1]. In biofilms, bacteria exhibit increased resistance to chemical, mechanical, and biological removal, complicating cleaning processes essential for preventing nosocomial infections, ensuring food safety, or maintaining efficient industrial operations [2, 3]. A particular attention has thus been devoted over the past two decades to the identification of the determinants of bacterial adhesion to solid surfaces as a means to design more efficient anti-fouling surfaces [4]. While chemical signals and biochemical interactions have been well-documented as regulators of adhesion [5–7], recent research highlights the influence of mechanical cues - such as surface softness and roughness — on bacterial colonization [8–14]. In addition, bacterial attachment often occurs in flow environments, where hydrodynamical forces are known to affect not only the rate of attachment and residence time, but also bacterial motility on surfaces [15–17].

However, despite immense efforts towards the development of efficient long-term anti-fouling surfaces, the fundamental mechanisms by which mechanical, chemical, and hydrodynamic factors (and their combinations) influence surface exploration, either through physical tuning of the interactions with surfaces or through sensing and translation, is still only partly explored [18]. This gap in knowledge is in part due to the absence of platforms that can independently and robustly manipulate physico-chemical properties of surfaces while also controlling nutrient and oxygen flow. Furthermore, most studies have focused solely on bacterial adhesion, neglecting the early stages of active surface exploration.

In the case of the common pathogen *Pseudomonas aeruginosa* (PA), we have recently observed a coupling between surface motility and microcolony formation — processes that typically take place within hours of attachment and are crucial for biofilm initiation [10]. In the present study, we present novel experimental approaches that allow for precise control over the physico-chemical and mechanical microenvironments at solid/liquid interfaces, coupled with time-resolved imaging of surface contamination. We demonstrate how these platforms can be leveraged to study early colony formation, advancing our fundamental understanding of bacterial surface exploration. Additionally, we propose a framework for controlling simultaneously the properties of solid/liquid interfaces and microfluidic flow, using PA as a model system to illustrate the potential of these techniques for investigating surface exploration dynamics, with a focus on the role of surface elasticity, chemistry and topography.

## 2 Material and methods

This section lists the chemicals and consumables used, and briefly describes the protocol for cell culture and the methods employed for data analysis and surface characterization. For the sake of clarity, the protocols for producing well-controlled surfaces and inclusion in fluidic devices are detailed in Appendices A-D. We have also included therein comments to stress important aspects of the sample preparation that we have found critical to achieve high-quality devices, but could otherwise be easily overlooked.

### 2.1 Reagents and consumables

Hydrogen peroxide (H_2_O_2_), sulfuric acid (H_2_SO_4_), acetic acid, acrylamide (AA, 40% in water), N,N’-methylenebisacrylamide (bisAA, 2% in water), ammonium persulfate (APS, ≥98%), N,N,N’,N’-tetramethylethylenediamine (TEMED, ≥99%), N(3-aminopropyl)-methacrylamide (APMA), 2-Hydroxy-4’-(hydroxyethoxy)-2-methylpropiophenone (Irgacure 2959, ref. 410896), 3-aminopropyltriethoxysilane (APTES), 1-ethyl-3-(−3-dimethylaminopropyl) carbodiimide (EDC), phosphate buffer saline without calcium and magnesium (PBS), glutaraldehyde (GTA), biotinylated fluorescein isothiocyanate (b-FITC), 2-(N-Morpholino)ethanesulfonic acid (MES, ref. M3671), Lyophilized streptavidin (SAv, ref. S4762), Bind-silane, and sigmacote were all purchased from Sigma-Aldrich (Saint Quentin Fallavier, France).

Ethanol, acetone and n-octane (all ≥99%), hydrochloric acid (37%), octadecyltriethoxysilane (OTE, 98%), 5/6-carboxyfluorescein succinimidyl ester (NHS-FITC, ref. 46410) and sodium borate buffer (pH 8.5, ref.28384) were purchased from Fisher Scientific (Illkirch, France).

NOA 85 and NOA 68TH optical glues were purchased from Edmund Optics (Lyon, France).

Lyophilized 1,2-dioleoyl-sn-glycero-3-phosphocholine (DOPC) and 1,2-dioleoyl-sn-glycero-3-phosphoethanolamine-N-(Cap Biotinyl) (DOPE-cap-b) were purchased from Avanti Polar Lipids (Alabaster, USA).

HEPES buffered saline (HBS; 10 mM HEPES, pH 7.4, 150 mM NaCl, all from Sigma) was prepared in ultrapure (UP) water (resistivity 18.2 MΩ.cm). Tryptone broth (TB) was purchased from Euromedex (Souffelweyersheim, France) and diluted in UP water to a concentration of 10 g.L^−1^.

Carboxyl beads were purchased from Polysciences (Hirschberg, Germany; Polybead Carboxylate Sampler Kit I, ref. 19819, and Polybead Carboxylate Microspheres 0.05*µ*m, ref. 15913).

IBIDI hard-plastic flow chambers and their connectors were purchased from IBIDI (Gräfelfing, Germany; sticky-Slide I Luer, ref. 80168, and Elbow Luer Connector, ref. 10802).

Polydimethylsiloxane (PDMS) was purchased from Neyco (Vanves, France; Sylgard 184, ref. DC184-5.5) and used at a 10:1 PDMS:crosslinker ratio as recommended by the manufacturer.

PDMS tubing (Tygon ND 100-80, ref. AAD04103, and E-3603, ref. ACF00001, both Saint Gobain) and polytetrafluoroethylene (PTFE) tubing (Adtech Polymer Engineering, ref. BIOBLOCK/05) were purchased from Fisher.

### 2.2 Bacterial strains, preparation and channel seeding

The strain used in this study was *Pseudomonas aeruginosa* wild-type (WT) PAO1. Bacteria were inoculated in Luria-Bertani (LB) medium from glycerol stocks, and grown overnight at 37°C and 250 rpm in a shaking incubator. The next morning, 10 *µ*L of the stationary phase culture were diluted in 3 mL of LB medium and incubated in the same conditions for 3.5 hours, to reach mid-exponential phase (OD600 = 0.6-0.8). Bacteria were then diluted to OD600 = 0.005 in our working medium, TB (in Section 3.2.2 and 3.3 experiments), or TB:PBS (w/o calcium and magnesium) with a volume ratio of 1:2 in Section 3.1 and 3.2.1 experiments. In TB:PBS bacteria proliferated less rapidly, but results were qualitatively similar.

### 2.3 Imaging and image analysis

Fluorescence analysis of the SAv platform homogeneity and mobility, and phase contrast imaging of bacteria, were performed on a confocal microscope (SP8, Leica, for sections 3.2.1, 3.2.2 and 3.3) or a wide-field microscope (Zeiss Axio Observer 7, for section 3.1), both equipped with a fully motorized stage and a 37°C incubation chamber. Images were typically acquired at 1 frame*/*minute.

Image processing (registration of stacks and segmentation) was performed using home-written macros in FIJI incorporating plugins “MultiStackReg” [19] and “Weka Trainable Segmentation” [20].

Explored area was calculated as follows: for each temporal sequence of segmented images, ten successive images (corresponding to a time interval of 10 minutes) were combined with the “OR” operator using a sliding time window. The covered area was quantified by counting the number of pixels, normalized by that of the single central image in the 10-minute series. Substracting 1 (so as to ensure that the explored area is zero for immobile, non-growing bacteria), multiplying by the average area per bacteria and dividing by 10 yields the explored area per cell and per minute for each time point, averaged over 10 minutes and over all bacteria in the field of view. To reduce stochastic noise, averaging is then performed over the experiment duration (as indicated in the text).

Characteristic growth time was calculated as follows: for each segmented image in a temporal sequence, the covered area was quantified by counting the number of pixels and values were normalized by that of the first image, to obtain the normalized surface density of bacteria as a function of time. The values were then fitted to a growing exponential exp (*t/τ*), with *τ* the characteristic growth time.

Analysis of the nearest neighbour distribution of obstacles (experimental and numerical curves) and Student’s t test on mobility data were performed using Mathematica. Stars represent p*<*0.05 (*), p*<*0.01 (**) and p*<*0.001 (***).

### 2.4 Contact angle measurements

Contact angles were measured on a home-built goniometer made of a vertical 10 *µ*L syringe used to bring water droplets in contact with the horizontal substrates. The shape of the backlit sessile droplets was imaged from the side using a 12-bit monochrome CCD camera (Chameleon USB2, 2560×1920 pixels, Point Grey Research) fitted with a macro zoom lens yielding a field of view of 11.14 × 8.36 mm^2^. Steady state advancing (resp. receding) angle was determined upon inflating (resp. deflating) the sessile drops and by image analysis using the “drop analysis” plugin of Fiji [21].

### 2.5 AFM elasticity measurements

The viscoelastic properties of the gels were characterized by AFM in 2 different ways, which provided consistent results:

1. Microrheology, as described previously [10]. Briefly, we used the “contact force modulation” technique [22] using a Nanowizard II AFM (JPK Instruments), with pyramidal-tipped MLCT probes (Bruker) of spring constant 15 mN/m. Data were analyzed using a home-written software for microrheology. A complete list of hydrogel composition, their elastic properties and an assessment of their spatial homogeneity can be found in [10].
2. Nanoindentation measurements with micrometer-size tips: We used a NanoWizard 4 Bioscience AFM (JPK instruments) to perform indentation measurements, and cantilevers with 1 *µ*m radius spherical tip (SAA-SPA-1UM, Bruker), spring constant 0.25 N/m and frequency 40 kHz. The indentations were performed in 8 × 8 force mapping mode at a size of 100 *µ*m (or 10 *µ*m for gradient indentation). For force-distance measurements, the indentation was performed at a speed of 2 *µ*m/s at a setpoint of 1 nN and a baseline of 3 *µ*m. To obtain the Young’s moduli, the Hertz model for a spherical tip was fitted to the measured force-distance curves using the proprietary JPK data analysis software. With the same software from the retraction curve we also obtained the adhesion force, the separation energy, and the adhesion length.

## 3 Results

One of the challenges that has to be addressed to study early surface exploration and colonization in the 1-10h time range is to simultaneously i) control a number of parameters defining the solid interface while ii) controlling shear and flow rate and ensuring sufficient renewal of nutrients and oxygen over the whole duration of the experiment in a setup that iii) permits sub-micrometric imaging of bacterial cells. This sets constraints on the flow device, whose dimensions should be small enough to ensure that the amount of medium required for the experiment stays reasonable while its renewal ensures enough supply for the growing number of bacteria in the channel. Microfluidic is therefore preferable to larger flow devices, raising in turn the challenge of circuit sealing with surface functionalization. In the following, we will demonstrate several realizations of devices allowing to study quantitatively the impact on surface colonization of different surface parameters.

We will focus on the example case of surface motility to illustrate the different approaches permitting accurate control of the above-mentioned environmental cues. In the model pathogen PA, surface motility is indeed a key parameter in exploration and colonization of the solid/liquid interface. The micrometer-long, contractile type-IV pili (T4P) that mediates this so-called twitching motility promiscuously bind free or surface-attached molecules, and then retract by depolymerisation of subunits at their anchor base on the cell membrane, applying forces in the 50-100 pN range [23, 24] and dragging the cell body across the solid surface. Pili have also emerged as a key tool that individuals use to mechanically probe and respond to their environment [25–27]. Studying how interface properties modulate twitching motility can thus provide new insight into the physical aspects of these sensing and regulation mechanisms.

### 3.1 Surface rigidity

When swimming bacteria encounter a submerged surface, they frequently transition to a surface-adhered state, which will next lead to the emergence of a structured biofilm via complex mechanisms that combine proliferation and aggregation. During the past decade, among the many parameters that influence this process, mechanical signals have appeared to be fundamentally important [28, 29], specifically forces modulated by substrate rigidity [9, 10, 12]. Several studies have demonstrated, in particular, that long-term biofilm development was modified on softer or stiffer substrates, both in terms of quantity and spatial organization of cells, with results that were strongly dependent on microorganisms [30–32]. Elasticity is thus a parameter to consider e.g. when trying to decipher the behaviour of promiscuous pathogens that contaminate different parts of the body, or when designing indwelling devices which are often the source of nosocomial infections. However, it still remains elusive how bacteria can integrate surface stiffness immediately upon contact and how this can influence the bacterial outcome. By directly modifying physico-chemical interactions between the bacterial body and the substrate, surface rigidity may first impact bacterial attachment rate [33]. After permanent attachment, surface colonization involves dynamical processes based on proliferation and motility, that can all be impacted by the deformability of the underlying substrate.

In order to systematically study the impact of substrate stiffness on bacterial behaviour, one difficulty lies in decoupling the rigidity from chemical properties. The use of hydrogels is an interesting option: it permits scanning a wide range of Young’s moduli without changing the chemical nature of the surface, by modulating the cross-linking ratio. Polyacrylamide (PAA) in particular is a highly biocompatible hydrogel, and provides access to rigidities from *<* 1 kPa to hundreds of kPa, similar to those of biological tissues. To systematically investigate the role of rigidity in early bacterial behavior on surfaces, we have designed thin, surface-bound, transparent PAA patches allowing for in situ imaging of adhering bacteria on an inverted microscope (fig. 1(a)). These substrates can be characterized by AFM nanoindentation, giving elastic moduli spanning over 2 orders of magnitude, while hydration of the gels changed marginally (within the range 91 - 98%), thus guaranteeing similar surface chemistry.

**Fig. 1.**
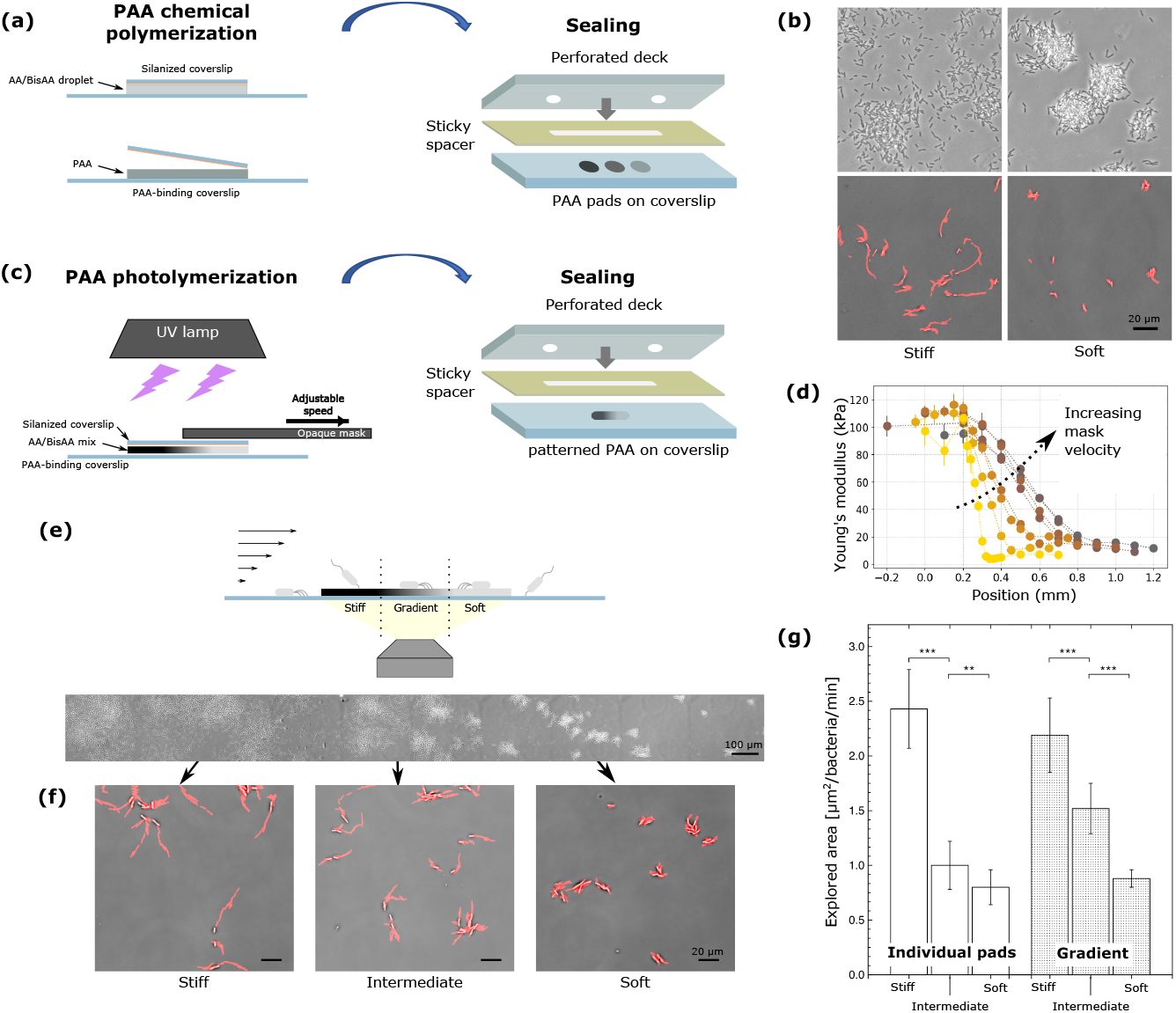
Creating substrates with varying rigidity for in situ observation of bacterial colonization. (a), PAA pads with controlled homogeneous rigidity are created via chemical polymerisation of an AA/bisAA mix, and can next be included in a sealed microfluidic device. (b), top: bacterial colony phenotype after ∼ 5h strongly depends on substrate elasticity, bottom: snapshots of bacteria on stiff and soft PAA after 30 min of experiment, and projected explored area over the following 30 min (here stiff ∼ 84 kPa, soft ∼ 3 kPa). (c), using photopolymerisation and a sliding opaque mask to modulate light exposure allows the spatial patterning of a single PAA pad. The resulting pad is, again, included in a flow device. (d), Young’s modulus measurements using AFM nanoindentation show linear stiffness gradients, with a slope that directly depends on the mask velocity. (e), experimental setup for phase contrast imaging of bacteria adhering to a PAA linear gradient under controlled flow, and example of heterogeneous bacterial colonization after 4.5h under constant shear rate (3.5 s^−1^). (f), snapshots of bacteria on stiff, intermediate and soft regions after 1.5h, and projected explored area over the following 30 min (stiffness values correspond to the ones measured on (d) for the dark-brown plot). (g), explored area per unit of time and per bacterium for the same three regions (right), averaged over the first 150 min, and for separate individual pads (left) of rigidity ∼84 kPa, ∼20 kPa and ∼3 kPa.

Inclusion of such substrates in a microfluidic device is however a challenge: indeed, plasma-sealing traditionally used to form PDMS-based microfluidic channels would damage hydrogel surfaces. Previous studies, however, have shown that hydrogels retained their mechanical properties after drying and rehydration (see [10], Supplementary Fig. 13(c)). We have thus developed an alternative approach in which hydrogels are first prepared and dried on a glass slide, onto which a flow cell is assembled via the use of double-sided sticky tape and a flat deck (figure 1(a)). This protocol allows the simultaneous imaging of bacteria on surfaces that are chemically similar but have different Young’s moduli in a single experiment, in the same channel, which limits experimental variations.

Using this approach, we have been able to highlight the strong impact of substrate elasticity on PA behaviour right after attachment. For a few hours after attachment, bacteria explore the surface by twitching motility, which strongly depends on substrate elasticity as shown on fig. 1(b) - see Subsection 2.3 for details on explored area calculation. In [10], to rationalize these results, we have developed a minimal model that shows that the net cell displacement across the substrate when a pilus contracts decreases on a soft deformable substrate, without assuming T4P attachment rate to significantly depend on rigidity, nor adhesion of the bacterial body that is mediated by exopolysaccharides. As colonization progresses, this twitching efficiency modulation strongly impacts bacterial 3D organization into microcolonies on the surface (fig. 1(b)).

Building on these earlier results, we have here employed photopolymerization to spatially pattern the rigidity of a given hydrogel susbtrate (fig.1(c)) [34]: using a photoinitiator and modulating exposure time using a sliding opaque mask allows the spatial patterning of gel crosslinking within a single PAA pad (see Appendix A for details). This permits, for example, the creation of stiffness gradients, i.e. regions of locally varying stiffness between a stiff and a soft homogeneous regions (fig. 1(d)). In that case, we can thus image *in situ* the behavior of bacteria on a heterogeneous substrate, similar to what mostly happens in practical cases, when they encounter for instance complex tissues. Fig. 1(e) illustrates such an example where bacterial behavior varies over a few hundred microns, and dense colonies coexist with neighboring dispersed progenies. Tracking bacteria highlights their difference in surface motility: the stiffer the substrate, the more persistent are individual trajectories (fig. 1(f)). This results in a clear increase in the explored area per bacterium per unit time with increasing substrate rigidity (fig. 1(g)). We reckon that this propensity to explore the underlying substrate is at the heart of PA strategy to spatially organize into microcolonies on surfaces.

### 3.2 Surface chemistry

#### 3.2.1 Ex situ surface functionalization: controlling surface polarity

The chemistry of the solid/liquid interface is another parameter of importance for the interactions between bacteria and a substrate. While some species bind specifically to well-defined chemical groups, a number of commensal bacteria can attach to most surfaces through interactions that are at present not fully understood [35]. How such organisms modulate or maintain adhesion onto chemically diverse substrates is central to the design of anti-fouling surfaces. Our model bacteria *P. aeruginosa* forms non-specific interactions both with its cell body and through its type-IV pili [24]: while pilus attachment occurs through direct interaction with the substrate, the cell also decorates the surface with extracellular matrix. This mix of macromolecules including a large fraction of exopolysaccharides mediates adhesion of the cell body and may also mediate Pilus attachment to a lesser extent [36].

In a model study to monitor how the chemistry of abiotic surfaces influences surface exploration by *P. aeruginosa*, we seeked to control surface charges and monitor how hydrophilic and hydrophobic interfaces compare in terms of environment exploration by surface-attached cells. To this aim, we have used silanes with long hydrophobic alkyl chains that can be densely grafted onto a glass coverslip: octyltriethoxysilane (OTE), a trivalent silane bearing a 18-carbon alkyl chain, self-organizes on the surface as a compact monolayer [37]; alternatively, sigmacote forms an hydrophobic coating through the binding of 1,7-dichloro-octamethyltetrasiloxane [38], an oligosiloxane terminated at both ends by a chlorosilane group. Both molecules expose similar methyl groups, but the molecular organization of the layers is different (self-assembled monolayer for OTE, disorganized structure of “loops” on the surface for sigmacote), allowing to disentangle hydrophobic vs supramolecular organization effects. In both cases, the contact angle of water droplets is greater than 100°.

Because of their hydrophobic nature, however, these reagents are grafted in organic solvents that are not compatible with the use of PDMS-or hard-plastic-based microfluidics. Functionalization is therefore performed before sealing of the channel, and plasma sealing - the most common method for sealing PDMS microfluidic channels - cannot be used because it would alter the chemistry of the functionalized surface. As an alternative, we have optimized a sealing method introduced by Agostini et al. [39] based on the use of UV-curing optical glue. The principle is illustrated on fig. 2(a): briefly, the glue is spin-coated onto a clean glass coverslip, and transferred onto the part of the PDMS that will be sealed onto the functionalized coverslip using stamping with minimal applied pressure. It is then carefully lifted to avoid spreading glue within the channels, deposited onto the dried functionalized coverslip, and subsequently sealed with UV light. We have systematically screened low-viscosity optical glues that permit depositing a thin glue layer onto the PDMS device while maintaining strong adhesion, such that this optimized protocol ensures resistance of the device to positive pressures up to a minimum of 900 mbar on a variety of surfaces (glass, sigmacote, APTES-coated glass, gold) while minimizing spreading within the functionalized channel: staining of the glue with NHS-FITC (diluted at 1 mg*/*mL in DMSO then at 10 *µ*g*/*mL in 50 mM sodium borate buffer, and incubated within the channel for 1h before rinsing) indicates that spreading does not exceed 1-2 µm within the channel, and that channels as small as 10 µm (width) x 5 µm (height) could be sealed with this method without significant narrowing or blockage (fig. 2(b,c)). A step-by-step protocol for robust and precise sealing is detailed in Appendix C.

**Fig. 2.**
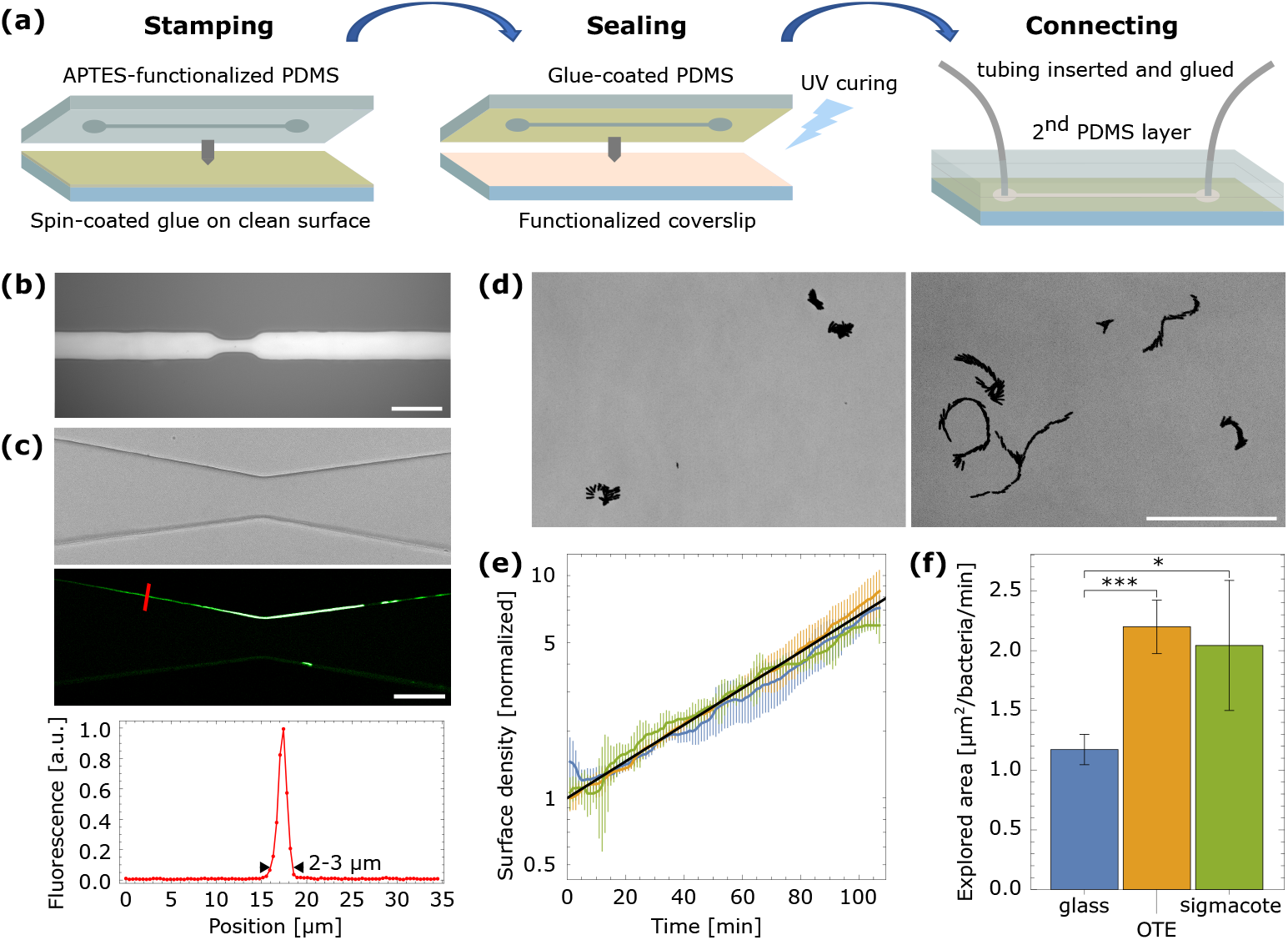
Controlled surface polarity in microfluidics and its effect on bacterial motility. (a), principle of inclusion of a functionalized coverslip into a PDMS-based microfluidic device using optical glue. (b), reflection image of a 15-*µ*m-thick channel with a 10-*µ*m-wide constriction sealed by gluing an APTES-functionalized coverslip. The reflection image demonstrates minimal spreading of the glue inside the channel. (c), phase contrast (top) and fluorescence confocal (middle) images of a glued channel with glue staining using NHS-FITC. Bottom, fluorescence intensity profile along the red line on the fluorescence image. (d), simultaneous monitoring of *P. aeruginosa* exploration of plasma-cleaned glass (left) and OTE-coated (right) coverslips included in glued PDMS channels. A 30-min time projection of phase contrast images is shown, with darker areas marking the regions explored by the bacteria. (e), surface density of bacteria over time, normalized to its initial value, for plasma-cleaned glass (blue), OTE (yellow) and sigmacote (green) surfaces, shows identical exponential growth. Black line, exponential fit to the ensemble-averaged data. (f), explored area per unit of time and per bacteria for the same three surfaces (glass, n=8; OTE, n=3; sigmacote, n=2 independent experiments). All scale bars are 50*µ*m

Using this protocol for including functionalized surfaces and juxtaposing two coverslips, we could image in parallel bacteria exploring hydrophilic and hydrophobic surfaces (fig. 2(d)). This quasi-simultaneous monitoring (alternating between the two surfaces within 1 minute) is key to quantitatively compare the two conditions. With this setup, we found that during the initial stages of surface colonization, the growth of bacteria is comparable on all surfaces, with a characteristic growth time 36.8 ± 0.9 min on glass, 34.7 ± 3.0 min on OTE and 36.9 ± 5.5 min on sigmacote (see Material and Methods for characteristic growth time calculation, and fig. 2(e) illustrating exponential growth). In contrast, motility is significantly affected: the area explored on hydrophobic surfaces is roughly doubled during the first 30 minutes on the surface compared to clean glass (fig. 2(f)).

This is in agreement with studies showing increased pilus adhesion but comparable cell body adhesion on hydrophobic vs hydrophilic substrates [24]. It does not match other observations using either PDMS as a hydrophobic substrate [40] or charged polymer increasing hydrophilicity [41], but these two studies were performed on bacteria confined in the interstitial space between an agar slab and the plastic surface of a Petri dish, observed without temporal monitoring after several hours: the observed discrepancies illustrate the importance of normalizing motility assays to draw conclusions that can be very different in nature. Indeed, the short-term monitoring presented in this paper mostly probes the physical impact of surface polarity on the twitching process (e.g., passive changes in cell body friction or in the strength of pilus attachment), while long-term assays on colonies capture a wealth of sensing and transduction phenomena that is expected to be central to the behaviour of bacteria. Another study has failed to find a correlation between surface polarity and twitching, but the experimental platform typically coupled varying rigidities, polarities and roughness in a non trivial manner, such that the effect of polarity was not probed independently [42], illustrating the importance of well-controlled platforms in which all relevant parameters (including shear forces and nutrient flow) are precisely controlled.

#### 3.2.2 In situ surface functionalization: towards probing interactions with complex biomolecules

While studying colonization of abiotic surfaces may shed light on contamination of medical devices or other artificial surfaces, or inform on the fundamental physical understanding of the twitching process, it is of key importance to study how pathogens interact with biotic surfaces. Biomimetic substrates offer the possibility to specifically decipher the interaction of pathogens with chosen components of cell surfaces, such as transmembrane proteins or polysaccharides that surround eukaryotic and prokariotic cells and mediate their interactions with other cells and extracellular matrices. Recently, such a synthetic model of eukaryotic cell membrane was used to study how the binding of PA lectin A to glycosphingolipid globotriaosylceramide (Gb3) could alter existing lipid domains [43, 44]. Of note, these studies did not use fluid flow and observation of bacteria was therefore restricted to short term imaging (*<* 1 h).

Contrary to more simple chemical functionalizations, indeed, biomimetic platforms incorporating complex biomolecules are challenging to include in microfluidic circuits. They are often deteriorated upon drying and require that an aqueous environment be continuously preserved after surface functionalization. As a result, the previously-described sample preparation strategy to incorporate such surfaces into a flow chamber, which relied on drying of the surface functionalization, must be adapted. In order to retain full flexibility on the microfluidic channel design, we have developed a protocol allowing surface biofunctionalization within a sealed PDMS channel while maintaining the quality of the resulting surface - a point that is rarely discussed in the literature but challenging when using PDMS channels from which free chains easily diffuse and contaminate the surface. The choice of this approach is dictated by the need to reduce functionalization volumes to a minimum when using biomolecules that can be either costly or only purified in small amounts: indeed, it is commonly estimated that full concentration equilibrium is reached after flushing at least 6 times the volume of the device (inlet tubing + microfluidic channel). A particular effort was thus devoted to minimizing the dead volume on the channel input side, which can be reduced to ∼ 6 *µ*L in our setup.

In addition, because bacteria can adhere to sub-micrometic defects in the functionalized layer, it is of utmost importance to minimize the presence of such defects, which critically depend on the contamination of the surface. By careful optimization of the functionalization process, we have found that key to the quality of the resulting layer are (i), appropriate preparation of the PDMS chip ensuring minimal contamination of the surface by remaining free PDMS chains; (ii), the use of tubing material minimizing biomolecule adsorption, which is significant for classically-used silicone tubing; (iii), appropriate rinsing and functionalization volumes that are estimated by taking into account the progressive removal of the initial solution in Poiseuille flows that is slowed down close to the walls (resulting in a factor ∼ 6 to achieve proper rinsing, see above); (iv), the careful tuning of flow rates to optimize functionalization times while avoiding spurious effects such as shear-rate-induced surface gradients of biomolecules [45]. A detailed protocol is presented in Appendix D.

Using this protocol, we could demonstrate high quality surface functionalization within microfluidic channels using a biotinylated supported lipid bilayer (SLB) onto which a partial monolayer of streptavidin (SAv) is anchored, providing binding sites for biotinylated molecules of choice onto which adhesion of bacteria can be studied (Fig. 3(a)). Here, as a proof-of-principle demonstration of the platform, we focused on the impact of the surface fluidity on bacterial motility at the onset of surface colonization. Indeed, SLB anchoring of SAv molecules provides a fluid surface with a typical diffusion coefficient around ≈ 1 *µ*m^2^*/*s, a scale relevant for PA pilus traction at velocities *v*_*c*_ ≈ 1 *µ*m*/*s over a few seconds. Alternatively, crosslinking of SAv molecules using glutaraldehyde (GTA) suppresses fluidity while maintaining the surface chemistry and its functionality (in particular biotin-binding ability) [46]. Homogeneity of the layers and diffusion coefficients with and without crosslinking were probed in situ after incubation with a biotinylated fluorophore (biotinylated fluoresceine isothiocyanate, b-FITC) using fluorescence confocal imaging and fluorescence recovery after photo-bleaching (FRAP) (Fig. 3(b) and (c)): in the absence of GTA crosslinking of SAv molecules, we measured a diffusion coefficient of *D* ≈ 0.9 *±* 0.1*µm*^2^*s*^−1^ and a mobile fraction *>* 99% (no immobile fraction could be detected from the fluorescent images when averaging over up to 20 images to reduce fluorescence shot noise); in contrast, for GTA-crosslinked layer, we found a mobile fraction *<* 1% (no mobile fraction could be detected from the fluorescent images when averaging over up to 20 images) and the diffusion coefficient was thus undefined.

**Fig. 3.**
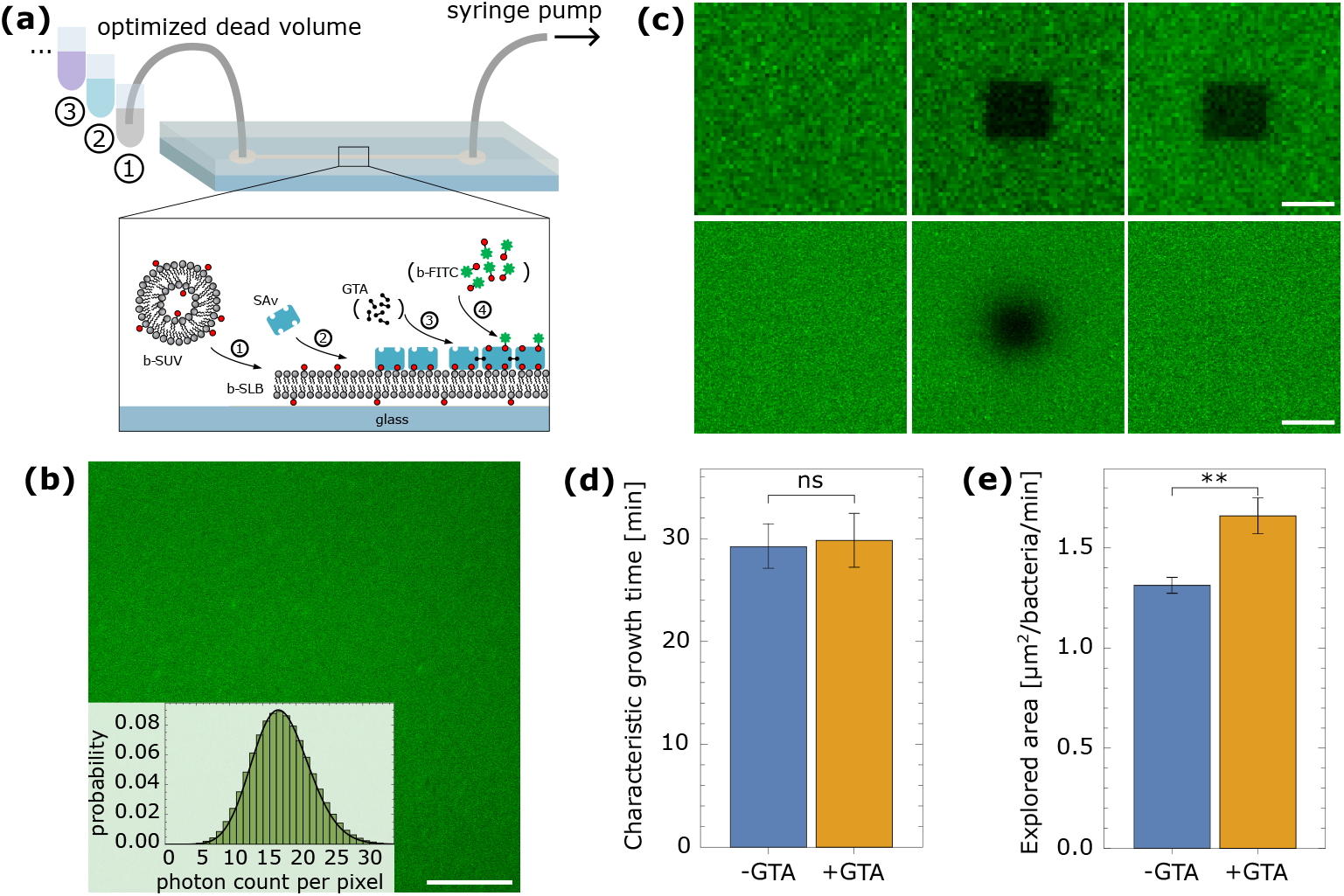
Complex surface biofunctionalization in a PDMS microfluidic chip and effect of fluidity on bacterial motility. (a), principle of surface biofunctionalization in a plasma-sealed PDMS microfluidic chip. (b), confocal fluorescence image of the functionalized surface labeled with b-FITC. The distribution of photon counts over the image matches the expected Poisson distribution of a spatially homogeneous sample. Scale bar, 20*µ*m. (c), snapshots of a FRAP assessment of the surface motility with (top) and without (bottom) GTA crosslinking. Images taken before (left), 5s after (middle) and 30 min after (right) bleaching over a 10 × 10*µ*m^2^ square area. Scale bar, 10*µ*m. (d), the characteristic growth time of the surface density of bacteria, for fluid (blue) and crosslinked (yellow) surfaces, demonstrating identical exponential growth. (e), explored area per unit of time and per bacteria for the same two surfaces (n=5 different fields of view per surface).

Taking advantage of the design flexibility offered by the use of PDMS microfluidic chips, we assessed the effect of surface fluidity on the mobility of bacteria using a chip containing two identical channels that were functionalized in parallel to create a SAv platform, one of them being further incubated with GTA. Comparing the characteristic growth time of bacteria on the two surfaces (fluid vs. immobile), we find that colonization rate is unaffected by the fluidity, indicating that it does not influence the ability to attach to the substrate (Fig. 3(d)). In contrast, mobility is significantly decreased on a fluid layer, indicating that twitching is less efficient when pili attach onto mobile biomolecules that do not provide fixed anchor points (Fig. 3(e)). The diameter of one pilus is ≈ 4 nm, meaning that on average each pilus will bind between a single and a couple hundreds SAv molecules, depending on the assumption that is made on its persistence length [23, 47] and hence on the length adhered on the surface: using Einstein-Smoluchowski relation, the drag coefficient, *ζ*, on a single SAv can be estimated from its diffusion coefficient *D* as *ζ* = *kT/D*, with *kT* the thermal energy. Using the average retraction speed *v*_*c*_ *≈* 1 *µ*m*/*s [48], the critical force required to drag the SAv molecules is around 4 fN (for one molecule) and up to 1 pN (for ≈ 250 molecules, corresponding to an adhered length of 1-2*µ*m), orders of magnitude below the retraction force experimentally measured for PA (≈ 30 pN [24]). Conversely, the cell body can create bonds with the underlying surface on a typical area ≈ 0.1 − 1*µ*m^2^, covering ≈ 10^3^ 10^4^ SAv molecules, inducing a drag in the range 5-100 pN on the fluid surface. As a result, a single pilus should not be able to drag the cell body by attaching to such a fluid layer, while 5-8 pili acting together (as frequently estimated for the PAO1 strain used in this experiment [25]) could just about equilibrate the drag on the cell body if the lowest part of the estimation range is considered. This hints that the observed cell motility on the fluid surface may rely partly on the presence of defects in the bilayer (allowing the pilus to stick firmly to the surface and drag the cell body), while adhesion to the fluid membrane might result in up to 50% of the mobility observed on the crosslinked platform. It is hence less efficient than on the latter. To further test the effect of surface fluidity, the use of larger biomolecules bound to SAv could permit screening the presence of such defects that should result in a more striking difference between fluid and crosslinked surfaces.

### 3.3 Surface topography

All the above-mentioned examples of surface functionalization result in flat surfaces. However, bacteria adhering on biotic or natural surfaces (e.g. cell layer, soil) encounter a much more complex topography with a nano- and micrometric scale rugosity which is known to impact the onset of surface colonization [14, 49, 50]. Efforts have mostly focused on adhesion rather than motility, and in particular on the effect of nanoscale roughness on contamination [51, 52]. Pioneering studies, however, have shown that surface discontinuities of height comparable to or larger than the size of the bacteria could act as obstacles for surface motility [53, 54]. While these studies have demonstrated that micron-scale roughness could significantly impact surface exploration, how random obstacles at controlled density, mimicking a more realistic rough surface, affect the colonization of surfaces remains to be deciphered.

Building upon our expertise in including well-controlled PAA hydrogels into microfluidic channels, we have developed a protocol to covalently graft polystyrene beads at controlled densities onto similar gels. To this aim, we use a mix of AA, bisAA and APMA to produce homogeneous gels with mechanical properties similar to that of classical PAA gels but providing surface-exposed amine moieties that can be used to couple carboxyl-functionalized polystyrene beads. To ensure that only the effect of topography is probed, and that we do not observe preferential attachment of bacteria to the beads or to the gel due to differences in surface chemistry, we use a mix of nanometric and micrometric beads at densities that permit covering entirely the surface of the gel. Nanometric beads form a uniform, dense layer with a roughness comparable to that of the raw gel (arithmetic roughness Ra ≈ 5-10 nm) that does not affect significantly the mechanical properties of the surface due to the absence of attractive inter-object interactions (as probed by AFM, see Methods and Table A1). Within this uniform layer, the density of micrometric beads coupled to the substrate is robustly determined by the fraction of microbeads in the initial bead mix added to the gel. Using a multiwell PDMS mask separating gels formed on the same coverslip permits creating surfaces with various roughnesses that can subsequently be incorporated in the same microfluidic channel.

As described in Appendix A, PAA gels can be dried for circuit sealing and subsequently rehydrated without damaging their surface or modifying their mechanical properties. This is however not the case of bead-functionalized gels that deform irreversibly around beads upon drying. In order to include these substrates into microfluidic devices without drying, we have thus used commercial devices based on a hard-plastic deck onto which a sticky tape defines the channel geometry. The benefit of this commercial solution is that it permits to robustly seal the circuit without fully drying the surface, with a channel long enough to allow probing of up to three gels with different topographies simultaneously.

This procedure can be used to couple obstacles of various sizes to the gel (Fig. 4(b)). For each of these sizes, we have characterized their distribution by computing the nearest neighbor distribution (NND) and comparing it to the case of a random distribution with excluded area (i.e., beads have a finite size and cannot overlap). An example for 2 *µ*m beads is shown on Fig. 4(c) for different bead densities, demonstrating that the spatial distribution of beads closely follows the expected result for a random distribution. This is in contrast with substrates used in most of the studies conducted on bacterial colonization of rough surfaces, for which roughness, lateral spatial scale and ordering are strongly interlinked (e.g., arrays of domes [55], pillars [56], nano- or microgrooves [8, 57]). Here, we can fully decouple the height for the obstacles from their density and mimic more realistically a disordered surface. A detailed description of the platform preparation is provided in Appendix B.

**Fig. 4.**
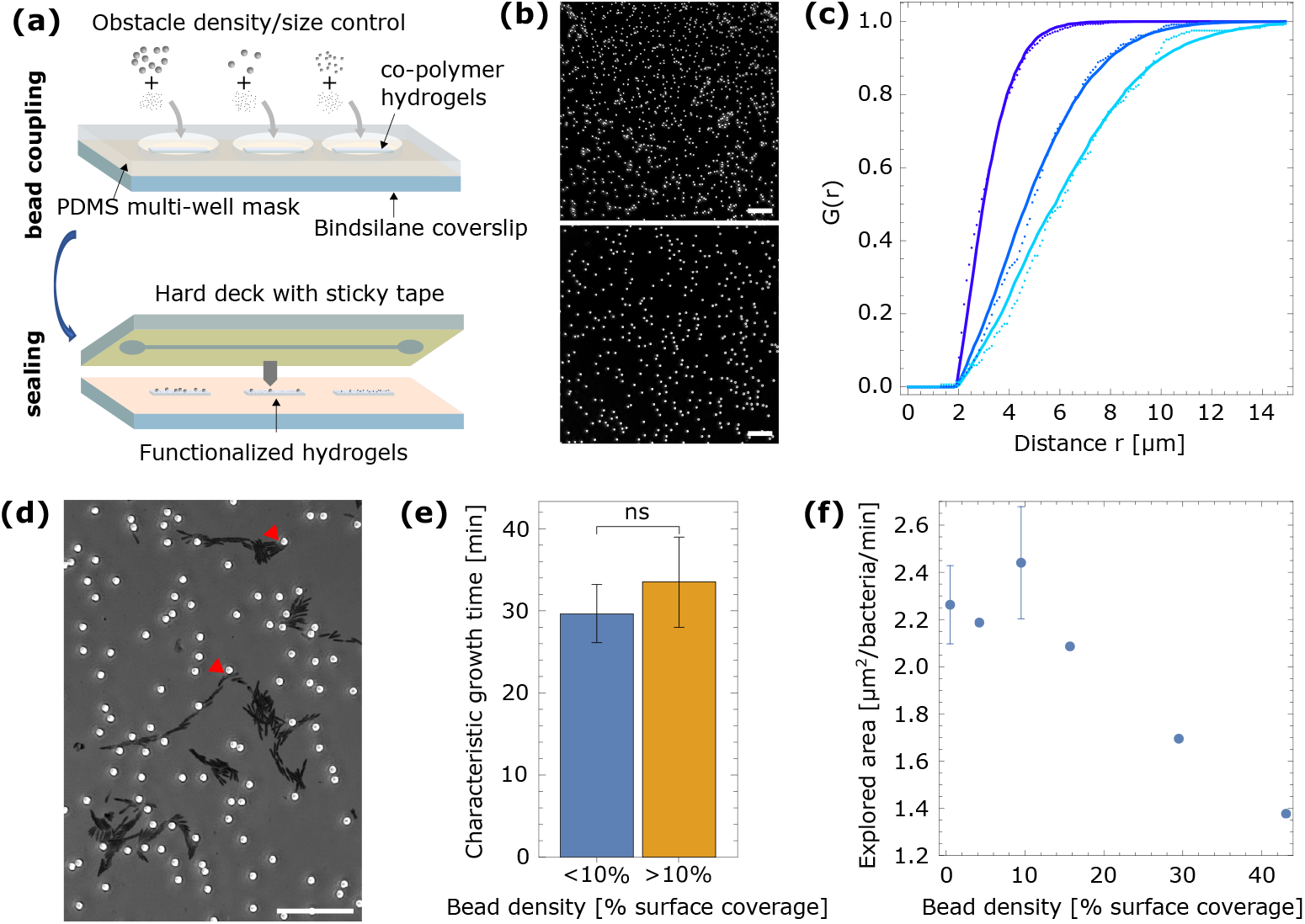
Controlled micrometric topography of surfaces and effect on bacterial motility. (a), principle of obstacle covalent binding and inclusion into a microfluidic device. (b), phase contrast imaging of gels functionalized with beads of 0.75 *µ*m (top) and 2 *µ*m (bottom) diameter. (c), nearest-neighbor distribution of 2-*µ*m beads for average surface densities of 11.2% (purple), 3.6% (blue) and 2.3% (light blue). These values refer to the percentage of area occupied by the beads. Dots are experimental values, lines are simulated distributions for the same average density and an exclusion radius around each bead equal to its diameter. (d), examples of trajectories of bacteria on a gel with 2-*µ*m beads at density 3%. A 30-min time projection of phase contrast images is shown, with darker areas marking the regions explored by the bacteria. Red arrows point to typical reorientation events upon encountering an obstacle. (e), characteristic growth time of the surface density of bacteria, for low (*<* 10%, blue) and high (*>* 10%, yellow) obstacle densities, demonstrating statistically equivalent exponential growth (n=3 experiments per condition). (f), explored area per unit of time and per bacteria as a function of obstacle density (n=2 fields of view per surface). Scale bars on all panels, 20*µ*m.

Following the previous subsections, we have demonstrated the interest of this platform on the example of surface exploration at the onset of colonization. Fig. 4(d) illustrates how 2 *µ*m beads act as obstacles that temporarily stop or reorientate the trajectory of bacteria, reducing the movement persistence. As a result, while the growth of bacterial coverage is unaffected (Fig. 4(e)), the area explored by bacteria at the onset of colonization (first 60 minutes) continuously decreases with the increase in the density of obstacles on the surface. This can be directly understood as an effect of the decrease in the processivity of bacterial twitching: while the instantaneous velocity between obstacles is mostly unaffected, the reduction in persistence length reduces the explored area by progressively transforming a short-time-scale ballistic displacement into a diffusive one (Fig. 4(f)).

## 4 Discussion

While an impressive number of studies have been published to decipher the key factors that influence antifouling surface design, well-controlled experimental platforms aimed at unraveling the fundamental mechanisms of bacterial exploration and colonization at solid/liquid interfaces remain relatively scarce in the literature (see Section 3 for examples). Such investigations critically depend on the robust and tunable integration of multiple experimental parameters, including: (i) the reproducible control of an ensemble of parameters of the solid surface, (ii) fluid flow conditions that impose hydrodynamic forces and medium renewal - and hence the availability of nutrients but also the ability to signal through secreted molecules — and (iii) time-resolved monitoring capabilities spanning from single-cell resolution to a large-scale view that allows direct comparison between surfaces. In this context, developing approaches that enable precise and reproducible tuning of individual surface characteristics is essential for gaining a deeper understanding of how the physico-chemical microenvironment influences the early stages of bacterial interaction and development. In this study, we used the example of PA short-term surface mobility to illustrate the utility of these integrated platforms. However, similar approaches can be applied to investigate adhesion alone or to study other bacterial species.

Fig. 5 recapitulates the key environmental parameters, and how model surfaces can be incorporated into microfluidic devices to ensure well-controlled experimental conditions. Indeed, constraints on the fluidic device vary depending on the type of surface functionalization (can the surface be functionalized in situ, or dried?), but also on the duration of the experiments (how much medium needs to pass through the circuit depending on its dimensions? Is gas permeability needed to maintain oxygen levels?). Importantly, meaningful data interpretation depends not only on careful tuning but also precise knowledge of the parameter(s) under investigation, and in situ characterization - as exemplified with surface fluidity measurements - can be a valuable asset for interpreting unexpected results.

**Fig. 5.**
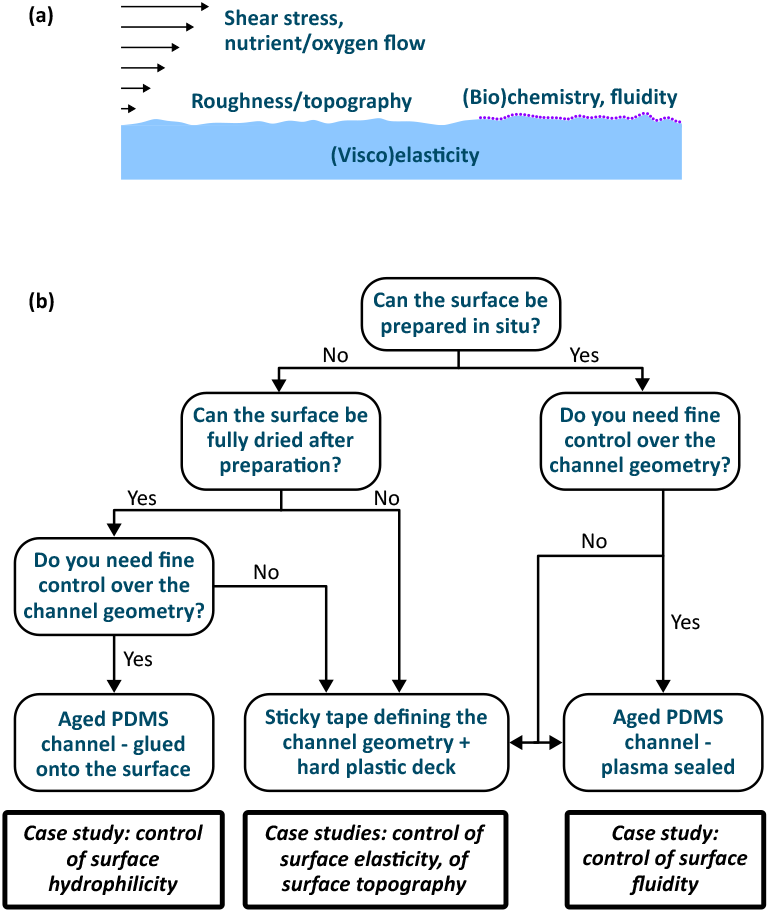
Summary of platform design. (a), physico-chemical cues at the solid/liquid interface. (b), decision tree for integrating functionalized surfaces into microfluidic devices. Case studies refer to the application examples demonstrated in the result section of this article.

Finally, while the surfaces discussed here represent the major properties currently known to influence bacterial adhesion, they should be seen as a foundation for further exploration. For instance, combining multiple parameters may help elucidate how bacteria integrate complex cues. Implementing spatial gradients at relevant biological scales—such as the size of individual cells, persistence length, or microcolonies—can shed light on taxis behaviors, as demonstrated in Subsection 3.1. Additionally, integrating more complex flow regimes (e.g., variable shear stresses, pulsatile flow, or spatial gradients in shear or nutrient composition) is feasible through protocols compatible with PDMS-based chips and soft lithography, offering promising avenues for future investigation.

## 5 Conclusion

In this paper, we presented a set of methods for integrating surfaces with well-defined physico-chemical properties into microfluidic devices, enabling the development of robust platforms to study how microenvironmental cues influence bacterial adhesion and motility during the initial hours of surface colonization. The detailed methodologies provided here aim to support the biophysics community in building upon pioneering studies that have begun to explore these complex interactions.

## Acknowledgments

The authors thank Ralf P Richter for fruitful discussions on SLB preparation, and Claude Verdier and Cendrine Moskalenko for help with the AFM experiments.

## Declarations

### Author contribution

M.L., K.C.E., R.M., Y.C.S., S.G.H. and E.M.V. designed experiments, and collected and analyzed data. L.B. designed experiments, supervised the project, and raised funds. S.L. and D.D. conceived the study, designed experiments, collected and analyzed data, performed numerical analyses, drafted the original manuscript, supervised the project, and raised funds. All authors contributed to manuscript review and editing.

### Data availability

Data sets used in this paper are available from the corresponding author on reasonable request.

## Appendix A Protocol - controlled elasticity platform

Samples preparation included two successive phases: (1) hydrogel pad preparation, (2) sealing in a microfluidic device.

1. **Gel preparation**: hydrogels of PAA were prepared following previously established protocols [58]. Rectangular glass coverslips (24×60 mm, no 1.5H, Menzel Gläser, Germany) were used as substrates for gel casting. They were piranha-cleaned (H_2_O_2_/H_2_SO_4_ = 1:3), plasma-cleaned in air for 3 min (Plasma surface cleaning system; Diener Electronic, Germany) and immediately immersed in a solution of Bind-silane (60 *µ*L of Bind-silane, 500 *µ*L of 10 % acetic acid, 14.5 mL of ethanol) for 1 hour before being rinsed with ethanol, water, and blow-dried with argon before use. Round glass coverslips (12 mm diameter, VWR, USA) were used as counter-surfaces for gel casting. After similar cleaning, they were immersed in sigmacote for 1 hour before successive 5-min sonications in acetone, ethanol and water, and blow-dried before use. Stock solutions of AA/bisAA were prepared in PBS and stored at 4°C until use. The final stiffness of the gels was tuned by adjusting the AA/bisAA content according to Table 1. PAA gels were obtained by adding 1 *µ*L of TEMED and 1 *µ*L of a freshly made APS solution (10 w% in water) to a volume of 168 *µ*L of AA/bisAA solution. A 3 *µ*L droplet of the mixture was immediately placed on the surface of a Bind-silane-treated glass coverslip, sandwiched by a sigmacote-treated round coverslip, and left for curing for 30 min in a water vapor-saturated atmosphere. Alternatively, to prepare stiffness gradients, the protocol was modified as follows [34]: 5 mg of Irgacure were added to 1 mL AA/bisAA premix. The solution was vortexed, and left at room temperature for 2h to ensure full dilution of the photoinitiator. 3 *µ*L of this solution were pipetted onto a bind-silane treated coverslip and flattened with a sigmacote-treated round coverslip. An opaque mask was stuck to the mobile part of a syringe pump (Harvard Apparatus, PicoPlus), and placed very close to the sample (≈ 1 mm), with its flat edge approximately at the center of the round coverslip. The setup was placed 1 cm below the surface of a UV lamp (Vilber VL-6.LC, 365 nm, 6W). The total exposure time was 2.5 min for all hydrogels. Once the mask was in place, the motor was activated to maintain constant speed for 2 min, after which the mask was quickly removed and the UV kept on for 30 more seconds. Using the AA/bisAA premix corresponding to a fully cross-linked Young’s modulus of 112 kPa (Table A1), this yielded hydrogels with 3 regions: a soft part (≈ 15 kPa), a gradient, and a rigid part (≈ 110 kPa). The mask velocity determined the width and slope of the stiffness gradient; e.g. at a velocity of ≈ 5.8 *µ*m*/*s, we obtained a gradient of ≈ 124 Pa*/µ*m, and at 0.8 *µ*m*/*s, a gradient of ≈ 850 Pa*/µ*m. In terms of gel formation, moving from chemical to photochemical polymerization relies only on changing the initiator, which does not influence the gel chemistry. Effects that could be potentially problematic are thus linked to the potential degradation of AA/bisAA by UV illumination. The photopolymerization of PAA, however, is a readily established approach that has been previously documented in the literature [34]. Previous studies have shown that surface properties of the hydrogel such as protein binding ability are preserved upon 365nm illumination. We have also checked that gel properties (rigidity, swelling – and hence hydration, binding of bacteria) were identical for illumination at 365 and 254 nm, and stable over at least 1 week, indicating that at the illumination doses used in this project, significant UV damage to PAA is unlikely as it would probably depend on the wavelength.
2. **Inclusion in a microfluidic device**: after curing, the round, sigmacote-treated coverslip was submerged in milliQ water, and lifted off using the tip of a scalpel blade, resulting in a circular pad of gel, of thickness ≈ 25 − 30 *µ*m, covalently bound to the bottom rectangular coverslip and exposing its free top surface. Circular gel pads were then scrapped with a razor blade in order to adjust their lateral size to the width of the microfluidic channels into which they would eventually be installed. Gel pads were then copiously rinsed with ultrapure water, and left for drying in a laminar flow cabinet. For gels with homogeneous mechanical properties, three such pads with different elastic properties, were prepared simultaneously on the same coverslip, arranged to fit along the length of the microfluidic channel. At this stage, the samples can be kept dry for several weeks without altering their mechanical properties or surface quality upon rehydration [10].

**Table A1.**
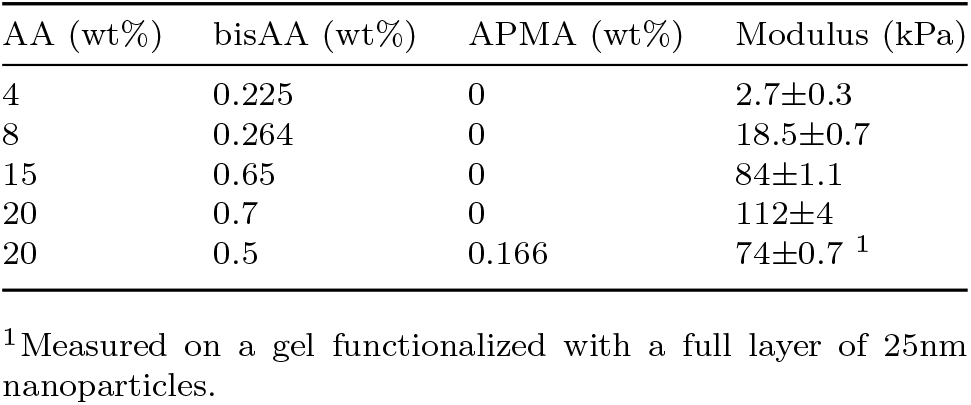
Hydrogel compositions and associated Young’s moduli measured by AFM.

| AA (wt%) | bisAA (wt%) | APMA (wt%) | Modulus (kPa) |
| --- | --- | --- | --- |
| 4 | 0.225 | 0 | $2.7 \pm 0.3$ |
| 8 | 0.264 | 0 | $18.5 \pm 0.7$ |
| 15 | 0.65 | 0 | $84 \pm 1.1$ |
| 20 | 0.7 | 0 | $112 \pm 4$ |
| 20 | 0.5 | 0.166 | $74 \pm 0.7$ <sup>1</sup> |

<sup>1</sup>Measured on a gel functionalized with a full layer of 25nm nanoparticles.

## Appendix B Protocol - controlled topography platform

Sample preparation included three successive phases: (1) hydrogel pad preparation, coupling of beads acting as obstacles, (3) sealing in a microfluidic device.

1. **Gel preparation** was similar to that of non-functionalized hydrogels, except that gels were prepared using mixes of AA, bisAA and APMA. The amount of added APMA was adjusted to provide one amine moiety per 300 AA monomeres (see Table 1). Three 12-mm-diameter 70-kPa gels were prepared at edge-to-edge distances of 10mm and scrapped after curing to reach a width of 5mm, matching that of the future microfluidic channel.
2. **Bead covalent coupling**: A home-made PDMS mold of thickness 2 mm and bearing three 15-mm-diameter circular holes was then carefully placed upon the coverslip to create a water-tight well around each of the gels. In each of the wells, mixes of carboxyl beads were coupled to the gels by adding (i) 250*µ*L of a 1mM EDC solution in MES buffer (100mM in ultrapure water, pH=4.5), (ii) 30*µ*L of a 1% v/v 50-nm-diameter carboxyl bead solution, and (iii) 0-250 *µ*L (depending on the aimed surface concentration of obstacles) of a 1% v/v solution of larger carboxyl beads (0.5 to 3 *µ*m diameter). Finally (iv), the volume was adjusted to a total of 600 *µ*L by adding the required amount of EDC solution, and the gels were incubated for 1h. During this time, the coverslip was kept in a closed petri dish to limit evaporation. After incubation, wells were rinsed with EDC using 5 times the incubation volume, and incubated further during 1h to ensure that all beads were fully coupled to the gel. Gels were subsequently rinsed again with ultrapure water using 5 times the incubation volume. The whole sample in the petri dish was then immersed in ultrapure water and the PDMS mold was carefully lifted off the coverslip without drying of the surface.
3. **Inclusion in a microfluidic device**: The coverslip was then removed from the water and partially dried by wiping water outside the gels until dry, while ensuring that the gels themselves were always kept hydrated. This permits to glue the IBIDI hard-plastic flow chamber around the gels, creating a microfluidic channel. This channel was immediately filled with ultrapure water to prevent drying of the gels. At this stage, the platform could be stored in the fridge for a few days before use.

To achieve a well-controlled density of obstacles with a high spatial homogeneity, as well as a good surface quality, it is essential to consider the following points: first, the beads should be gently mixed (e.g. with a pipette) at least once during incubation, to homogenize their distribution on the surface. Second, low-bind pipette tips were systematically used to prevent depletion of beads in the solution and accurate control of bead incubation concentrations. Finally, it is critical to ensure that the gels were never fully dried after curing, either when adhering the PDMS mold or when gluing the hard-plastic desk: indeed, we have found that drying results in irreversible deformations, possibly due to spurious bond formation between the beads and new regions of the gel.

## Appendix C Protocol - controlled surface polarity platform

Samples preparation included four successive phases: (1) surface functionalization, (2) PDMS microfluidic channel preparation, (3) sealing of the microfluidic device, (4) sealing of connectors for the tubing.

1. **Surface functionalization**: glass coverslips of 35 mm diameter (VWR, USA) were piranha cleaned and, for hydrophobic surfaces, functionalized using sigmacote (1h incubation, followed by rinsing with acetone, ethanol and water, and blow-dried before use) or OTE. For the latter, 40 *µ*L of OTE was added to 100 mL of n-octane in presence of 80 *µ*L of hydrochloric acid. The solution was stirred and cooled to 16°C for 15 min before being added to a beaker containing the coverslips in a teflon holder. Grafting was left to proceed for 60 minutes while maintaining stirring and cooling at 16°C, followed by thorough rinsing with acetone, ethanol and water, and drying in an Argon stream. Such a procedure was found to yield OTE monolayers of 2.5 ± 0.3 nm in thickness (as measured by ellipsometry on oxidized silicon wafers), compatible with the length of the fully stretched OTE molecule pointing upward on the surface. These coverslips can be kept either immersed in UP water (piranha-cleaned coverslip) and plasma treated before use, or dry (OTE and sigmacote) in a sealed box under neutral atmosphere, for several weeks.
2. **Microfluidic channel preparation**: channels 2 cm long, 100 *µ*m high and 500 *µ*m wide were molded in PDMS using soft lithography. Curing was performed at 135°C for 48 hours to ensure optimal crosslinking of the PDMS chains. The channel was then cut and detached from the mold, and entrance and exit were punched using a 1-mm biopsy punch. The prepared channels can be kept protected from dust in a sealed petri dish for several weeks.
3. **Gluing of the PDMS channel onto the functionalized coverslip**: this should be done shortly before using the device, to avoid potential contamination of the glass surface by airborne molecules or leftover free PDMS chains. A solution of 300 µL of APTES in 15mL UP water was left to hydrolyze on a magnetic stirrer for one hour in a glass beaker. During this time, on a 35-mm piranha-cleaned coverslip (different from the one that would be glued to the PDMS circuit), 500µL of NOA 85 optical glue were spin coated (10 min, 6000 rpm) to create a thin glue layer that was pre-cured for 40s using UV exposure (model BM 1200D, Chemical Electronique, France). At the end of APTES hydrolysis, the PDMS channel was exposed to plasma in air for 3 min (Plasma surface cleaning system; Diener Electronic, Germany) and immediately immersed in the filtered APTES solution (using a 0.2µm-pore-diameter syringe filter) for 5 min. It was then rinsed with UP water, dried with argon and placed onto the glue-covered coverslip. When depositing the channel, it is important that all parts outside the channel come in contact with the glue, but without pressing too strongly on the PDMS and avoiding slipping: this is essential to ensure that the glue will not spread within the channel itself. The PDMS was then lifted carefully from the coverslip and positioned on the functionalized coverslip (glass, OTE or sigmacote). It was pressed gently as required to remove air bubbles at the PMDS/glass interface, always avoiding slipping or excessive pressure. Finally, the circuit was sealed by curing the glue with UV for 20 min.
4. **Sealing of the tubing connectors**: As a last step, 3-cm-long stubs of PTFE tubing with an external diameter of 0.76 mm were inserted in the punched holes of the PDMS chip and fixed with NOA 68TH optical glue using an additional 20 min UV exposure. Lastly, an additional layer of PDMS was molded onto the circuit using a home-made circular mold clenched to the coverslip. The aim is to rigidify the device, strengthening adhesion to the coverslip. The additional layer also ensures better integration of the tubing stubs. About 2g of the PDMS-crosslinker mix were used to create a 28-mm-diameter device of about 5 mm height. PDMS was cured for 2h at 65°, and the channel was then used within a few hours. Of note, the external diameter of the PTFE tubing stub permits tight connection with the deformable Tygon tubing (0.02 inches inner diameter) by simple insertion.

To obtain well-controlled surface states and robust gluing of the circuit, we have found that the following points were critical:

- **Surface preparation**: full cleaning of the coverslips with piranha, followed by plasma activation in case the surfaces are kept between cleaning and functionalization, is essential to achieve homogeneous hydrophobic layers with a minimum of defects, as checked using contact angle measurements (OTE: advancing and receding contact angles of water *θ*_*A*_ = 112 ± 1° and *θ*_*R*_ = 98 ± 1°; Sigmacote: *θ*_*A*_ = 104 ± 1° and *θ*_*R*_ = 75 ± 1°).
- **PDMS preparation**: extended curing of PDMS at high temperature is essential both to minimize the presence of free chains that can diffuse on the surface and contaminate the glass surface, and to ensure efficient gluing: indeed, while plasma exposure can oxidize the PDMS surface and create -OH groups that can efficiently bind to the optical glue, turnover of chains at the air/PDMS interface quickly returns the surface to its initial state, and the characteristic time over which hydrophilicity is maintained critically depends on the presence of free chains resulting from incomplete crosslinking [59]. The curing parameters indicated here maximize this characteristic time while ensuring that the PDMS does not become brittle either due to excessive curing or to the use of an excess of crosslinker.
- **Gluing**: as mentioned above, the time window during which PDMS can bind efficiently to the glue after plasma exposure is short (typically 10-20 min). It is therefore essential that APTES functionalization be performed right after plasma exposure, and that glue stamping and curing be performed straightaway afterwards. Also, the choice of NOA85 glue accounts for its ability to bind to different surfaces and its low viscosity (ensuring that a thin layer can be deposited onto the PDMS). Conversely, NOA68TH was chosen to glue the tubing because of its high viscosity that prevents spreading through the punch holes and subsequent channel blocking. More than 10 alternative references have been tested to optimize the binding strength and limit spreading of glue within the channel, and we have found the reference provided above as the most suitable for this use.
- **UV curing**: full curing of the glue is essential both to maximize the strength of binding and avoid unwanted contamination of the functionalized surface. However, care must be taken to avoid damaging the functionalized surface in the process. Illumination around 365nm should thus be preferred to deep-UV illumination. In our experiments, we have verified that the contact angle for hydrophobic surfaces was unaffected by the curing process.
- **Tubing connection**: the two-step procedure for connecting the tubing to the microfluidic circuit stems from the observation that the appearance of leaks under pressure was always observed around the tubing insertion point due to extra mechanical constraints on the PDMS at these location. Using tubing with external diameter smaller than the punch size and gluing ensures that these mechanical constraints are minimized and that resistance to pressure of the final device is optimal.

## Appendix D Protocol - controlled surface biofunctionalization platform

Samples preparation included three successive phases: (1) Small unilamellar vesicles (SUVs) preparation, (2) PDMS microfluidic channel preparation and sealing, (3) surface functionalization.

1. **SUV preparation**: SUVs containing 5 mol % biotinylated lipids were prepared as previously described [60]. Briefly, lipids were dissolved in chloroform, mixed at a molar ratio of 95% DOPC and 5% DOPE-cap-b, and dried under a stream of nitrogen gas, followed by drying in a vacuum desiccator for at least 2 h. The lipid mixture was then resuspended in HBS at a final concentration of 2 mg/mL and homogenized by five cycles of freezing, thawing, and vortexing. The lipid suspension was sonicated with a tip sonicator (FB120; Fisher Scientific, UK) in pulsed mode [duty cycle: 1 s on (at 70% maximal power)*/*1 s off] with refrigeration for a total time of 30 min, followed by centrifugation (10 min at 13500 rpm) to remove titanium particle debris from the sonicator tip. SUVs were stored at 4 °C under an inert gas (nitrogen or argon) until use.
2. **Microfluidic device preparation**: channels 2 cm long, 100 *µ*m high and 500 *µ*m wide were molded in PDMS and cured as described above (48h, 135°C). 35-mm-diameter coverslips (VWR, USA) were cleaned with piranha and a PDMS microfluidic device including two parallel rectangular channels was sealed onto it with plasma after punching 0.5-mm-diameter holes for tubing insertion. After curing for 2h at 65°C the circuit was exposed to plasma again (in air, 3 min) before connecting the tubing (PTFE tubing, 0.76 mm external */* 0.3 mm internal diameter; inlet: 80 mm total length, corresponding to an inner volume of 5.6 *µ*L; outlet: about 1 m), and used immediately.
3. **Surface functionalization**: the outlet tubing was connected to a 10-mL glass syringe (gastight luer-lock syringe, SGE, ref. 008760) placed on a syringe pump (KDS Legato 210). 50 *µ*g*/*mL SUVs in HBS were injected at 10 *µ*L*/*min for 10 min and left to incubate for 30 min to form a supported lipid bilayer (SLB) by vesicle rupture and spreading. The circuits were then rinsed by injecting HBS at 10 *µ*L*/*min for 10 min. The SLB-coated surface was then incubated with 20 *µ*g*/*mL SAv in HBS for 60 min (10 min at 10 *µ*L*/*min followed by 50 min at 1 *µ*L*/*min) to form a dense SAv monolayer presenting biotin-binding sites, and excess SAv was again removed by washing with HBS as described above. One of the channels was then crosslinked using glutaraldehyde (8% in weight in HBS, 10 min at 10 *µ*L*/*min injection followed by rinsing as described above), and finally the surfaces were stained using biotinylated fluorescein isothiocyanate (b-FITC, 20 *µ*g*/*mL, 10 min at 10 *µ*L*/*min followed by 50 min at 1 *µ*L*/*min, then rinsing as described above).

To obtain well-controlled surfaces with a low density of defects, we have found that the following points were critical:

- **Surface preparation**: plasma exposure of the glass surface immediately before SUV injection is essential to the efficient spreading of vesicles creating a lipid bilayer with a minimum of defects. This additional plasma treatment should be performed before inserting the tubing to ensure that the surface inside the microfluidic channel is exposed to plasma. Of note, the PDMS surface is also activated and we found that a high-quality SLB is also created on the PDMS walls of the microfluidic channels.
- **PDMS preparation**: as in the case of ex situ surface functionalization (see above), extended curing of PDMS at high temperature is essential to minimize the presence of free chains that otherwise contaminate the glass surface and degrade the quality of SLBs.
- **Functionalization with b-FITC**: prior to injection, the fluorophore solution in HBS was deliberately partially photobleached to reduce the concentration of fluorescent b-FITC [46] to a level that essentially avoided self-quenching on the surface.
- **Tubing**: using PTFE tubing minimizes adsorption of biochemicals on the tubing walls compared to silicone tubing. It also reduces the elasticity of the circuit and hence the lag time when stopping the flow to exchange the inlet reservoirs: this is essential to avoid introducing bubbles in between functionalization steps.

